# Uniform cerebral organoid culture on a pillar plate by simple and reproducible spheroid transfer from an ultralow attachment well plate

**DOI:** 10.1101/2023.04.21.537886

**Authors:** Prabha Acharya, Pranav Joshi, Sunil Shrestha, Na Young Choi, Sehoon Jeong, Moo-Yeal Lee

## Abstract

Human induced pluripotent stem cell (iPSCs)-derived brain organoids have potential to recapitulate the earliest stages of brain development, serving as an effective *in vitro* model for studying both normal brain development and disorders. However, current brain organoid culture methods face several challenges, including low throughput, high variability in organoid generation, and time-consuming, multiple transfer and encapsulation of cells in hydrogels throughout the culture. These limitations hinder the widespread application of brain organoids including high-throughput assessment of compounds in clinical and industrial lab settings. In this study, we demonstrate a straightforward approach of generating multiple cerebral organoids from iPSCs on a pillar plate platform, eliminating the need for labor-intensive, multiple transfer and encapsulation steps to ensure the reproducible generation of cerebral organoids. We formed embryoid bodies (EBs) in an ultra-low attachment (ULA) 384-well plate and subsequently transferred them to the pillar plate containing Matrigel, using a straightforward sandwiching and inverting method. Each pillar on the pillar plate contains a single spheroid, and the success rate of spheroid transfer was in a range of 95 - 100%. By differentiating the EBs on the pillar plate, we achieved robust generation of cerebral organoids with a coefficient of variation (CV) below 19%. Notably, our spheroid transfer method in combination with the pillar plate allows miniaturized culture of cerebral organoids, alleviates the issue of organoid variability, and has potential to significantly enhance assay throughput by allowing *in situ* organoid assessment as compared to conventional organoid culture in 6-/24-well plates, petri dishes, and spinner flasks.

## Introduction

The human brain is one of the most complex organs in terms of its structural development and function (1–3). Studies on understanding the brain development and function have been carried out extensively over the past decades. Animal models have been widely used to understand the principle of human brain development. However, there are significant structural differences between humans and animals such as rodents (4,5). The lack of reliable *in vitro* models and the accessibility of human brain tissue have severely hindered the research on modeling brain development and disease (6). However, advancements in ability to differentiate pluripotent stem cells (PSCs) into three-dimensional (3D) organ-like tissue aggregates (a.k.a., organoids) have provided new opportunity to uncover development, structure, and function of the human brain *in vitro* (7). Organoids are 3D cell aggregates that resemble the native tissues or organs with similar cell type, function, architecture, and complexity, bridging the gap between traditional 2D cell culture models and animal models (8,9). Notably, brain organoids generated from induced pluripotent stem cells (iPSCs) not only mimic the cellular composition but also replicate the tissue structure and developmental aspects of the human brain. Hence, brain organoids hold a significant promise as a useful tool for investigating diverse aspects of human brain, including its development, functions, and brain-related disorders (7,10).

Several protocols, including guided and unguided methods, have been published for differentiating iPSCs into brain organoids. Unguided differentiation methods are used to generate whole brain organoids (11) whereas guided differentiation methods employ patterning factors to generate specific regions of the brain such as the hindbrain (12), midbrain (13–15), and forebrain (16,17). In unguided differentiation, embryoid bodies (EBs) were developed from iPSC aggregates, which are then embedded in extracellular matrices (ECMs) and cultured to promote neuroepithelial expansion and differentiation (11). The existing approaches of brain organoid generation involved several steps, including EB formation through aggregation in ultralow attachment (ULA) well plates, subsequent manual embedding of EBs in ECMs in 6-/24-well plates, and transferring these organoids to various culture platforms for further maturation, such as orbital shakers (1), spinner flasks (18), miniature bioreactors (16) or microfluidic devices (19). However, conventional organoid culture platforms such as 6-/24-well plates, petri dishes, and spinner flasks are cumbersome and costly for high-throughput applications (20,21) due to difficulty in *in situ* organoid imaging and a large volume of cell culture media and soluble factors required. In addition, these platforms face the issue of limited diffusion of nutrients and oxygen, resulting in necrotic cores within brain organoids and reduced functionality (22). To address these issues, microfluidic devices have been introduced in brain organoid culture, leading to enhanced organoid maturity and functionality (19,23). Nonetheless, microfluidic devices have technical challenges, such as low throughput (19) due to cumbersome steps of transferring organoids, requirement of user-unfriendly pumps and tubing, and incompatibility with high-throughput screening (HTS) equipment. These challenges need to be overcome to facilitate the widespread use of organoids for disease modeling and predictive compound screening.

In this study, we have developed a unique pillar plate and a complementary deep well plate to miniaturize brain organoid culture while enhancing throughput and reproducibility in organoid generation. In addition, we introduced a robust protocol for transferring and encapsulating EBs formed in an ULA 384-well plate onto the pillar plate containing Matrigel spots. This approach allowed the differentiation of multiple EBs into brain organoids individually and simultaneously on the pillar plate that can be used subsequently for *in situ* organoid imaging. In summary, the pillar/deep well plates, combined with the unique spheroid transfer protocol, represent a viable solution for the consistent generation of human brain organoids, which are crucial for applications of organoids such as high-throughput compound screening in clinical and industrial settings.

## Materials and Methods

### Design of the pillar/deep well plate for spheroid and organoid culture

The 36PillarPlate, 144PillarPlate, and 384PillarPlate were manufactured by injection molding of polystyrene and were functionalized with hydrophilic polymers (Bioprinting Laboratories Inc., Dallas, TX) (**Fig. 2**). The 36PillarPlate, 144PillarPlate, and 384PillarPlate contain 6 x 6, 12 x 12, and 16 x 24 array of pillars in 4.5 mm pillar-to-pillar distance, respectively, which allow to load up to 6 µL of cells suspended in hydrogel per pillar. The pillar plate with cells can be sandwiched onto the clear bottom 384DeepWellPlate (Bioprinting Laboratories Inc.) with cell culture media for static organoid culture. For cerebral organoid generation, the pillar plate was additionally coated with 2 µL of 1.5% (w/v) alginate in distilled water per pillar, completely dried, and then rehydrated before using by sandwiching the alginate-coated pillar plate onto the deep well plate with 20 µL of water at the well bottom. The alginate coating was introduced to prevent 2D cell growth on the pillar surface during long-term organoid culture.

### Human iPSCs culture and maintenance

Human iPSCs (EDi029A) were obtained from Cedar Sinai Biomanufacturing Center and maintained by following the manufacturer’s recommended protocol. Briefly, iPSCs were cultured in a 6-well plate coated with 0.5 mg Matrigel™ (354230, Fisher Scientific) in complete mTeSR plus media (100-0276, Stem Cell Technologies). The iPSCs were passaged every 4 - 5 days when it reached 80 - 90% confluency using the StemPro^®^ EZPassage™ disposable stem cell passaging tool (23181-010, Life Technologies). The cells were passaged as a small colony with a split ratio of 1:6 and cultured by changing the medium every day. iPSCs at the early passage number up to 10 - 12 stored in a frozen vial were suited for generating cerebral organoids.

### Formation of neural stem cell (NSC) spheroids

Human ReNcell VM (SCC008, MilliporeSigma) was cultured in ReNcell NSC maintenance medium (SCM005, MilliporeSigma) supplemented with 20 ng/mL basic fibroblast growth factor (bFGF, EMD Millipore), 20 ng/mL epidermal growth factor (EGF, EMD Millipore), and 1 % (v/v) penicillin/streptomycin (P0781, MilliporeSigma) with medium change every alternate day. Cells were passaged when they reached 90% confluency using Accutase^®^ (A6964, MilliporeSigma), resuspended in the complete maintenance medium, and centrifuged to remove supernatant. The cells were passaged as single cell suspension in the range of 1 - 1.5 million cells in a freshly coated T-75 flask with 20 µg/mL laminin (CC095, EMD Millipore). ReNcell was detached from the T-75 flask with Accutase, dissociated into single cells, and seeded in an ultralow attachment (ULA) 384-well plate (ULA-384U-020, Nexcelom) at a different seeding density of 500, 1000, 3000, and 9000 cells/well to form spheroids in the complete maintenance medium with medium change every alternate day. The spheroids were cultured in the ULA 384-well plate for 4 days.

### Formation of embryoid bodies (EBs) with iPSCs

iPSCs colonies were harvested with Accutase^®^ and dissociated into single cells using the complete human embryonic stem cell (hESC) medium consisting of DMEM/F12 (11320033, ThermoFisher), 5x knockout serum replacement (10828010, ThermoFisher), 100x GlutaMAX (35050-038, Invitrogen), 100x MEM-nonessential amino acids (MEM-NEAA; M7145, MilliporeSigma), 33x hESC quality FBS (10439001, ThermoFisher), 2-mercaptoethanol (21-985-023, Fisher Scientific) with a final concentration of 4 ng/mL basic fibroblast growth factor (bFGF; 100-18B, Peprotech) and rho kinase inhibitors such as a CEPT cocktail consisting of 5 µM Emricasan (50-136-5235, Fisher Scientific), 50 nM Chroman 1 (7163, Tocris), 0.7 µM Trans-ISRIB (5284, Tocris), and polyamine supplement diluted at 1:1000 (P8483, MilliporeSigma), or 10 µM Y-27632 (SCM075, Fisher Scientific). Single cells were suspended in the complete hESC medium containing either the CEPT cocktail or Y-27632, seeded in the ULA 384-well plate at a seeding density of 1000 cells/well, maintained in the hESC medium with CEPT or Y-27632 until day 4 of culture, and then cultured in the hESC medium without CEPT or Y-27632 for three additional days. The growth factor bFGF was used only on the day of cell seeding.

### Spheroid transfer to the 36PillarPlate from the ULA 384-well plate

Multiple parameters were optimized to establish the spheroid transfer method for rapid and robust encapsulation of EBs in Matrigel on the pillar plate. Briefly, four parameters were investigated, including (1) the size of spheroids transferred, (2) the time of spheroid transfer, (3) the concentration of Matrigel on the pillar plate, and (4) pre-gelation of Matrigel on the pillar plate prior to the pillar plate sandwiching onto the ULA 384-well plate. ReNcell spheroids cultured for 4 days at a seeding density of 500, 1000, 3000, and 9000 cells/well in the ULA 384-well plate were transferred to the alginate-coated pillar plate with 5 µL of 4 - 6 mg/mL of Matrigel by a simple sandwiching and inverting method. The pillar plate with Matrigel was sandwiched onto the ULA 384-well plate with spheroids, and then the sandwiched plates were inverted and incubated for 20 - 30 minutes at 37°C in a CO_2_ incubator for spheroid transfer and encapsulation in Matrigel. The spheroids on the pillar plate were cultured in the complete NSC medium in the 384DeepWellPlate. Similarly, EBs formed for 6 days at a seeding density of 1000 and 3000 cells/well in the ULA 384-well plate with the complete hESC medium were transferred to the pillar plate with 5 µL of 6 - 8 mg/mL of Matrigel by the sandwiching and inverting method. For robust EB transfer and encapsulation on the pillar plate with minimal dilution of Matrigel in the ULA 384-well plate, the pillar plate with Matrigel was pre-incubated at room temperature for 0, 5, and 10 minutes. After pre-incubation, the pillar plate with Matrigel was sandwiched onto the ULA 384-well plate with EBs, and then the sandwiched plates were inverted and incubated for 30 - 40 minutes at 37°C in the CO_2_ incubator for EB transfer and encapsulation. The EBs on the pillar plate were cultured in the neural induction medium (NIM) consisting of DMEM-F12, N2 supplement (17502001, ThermoFisher), GlutaMax, MEM-NEAA, and heparin (H3149, MilliporeSigma) in the 384DeepWellPlate for 5 - 6 days with medium change every alternate day.

### Differentiation and maturation of cerebral organoids on the 36PillarPlate

After 5 - 6 days of EB culture in the NIM to generate neuroectoderm (NE), NE on the pillar plate was differentiated in cerebral organoid differentiation and maturation medium without vitamin A (CDM-vit A) in the 384DeepWellPlate for 31 days with medium change every other day. The CDM-vit A consists of an equal volume of DMEM-F12 and neurobasal medium (21103049, Fisher Scientific) with N2 supplement, B27 supplement without vitamin A (12587010, ThermoFisher), insulin (I9278, MilliporeSigma), GlutaMax, MEM-NEAA, penicillin-streptomycin, and 2-mercaptoethanol.

### Differentiation and maturation of cerebral organoids in Matrigel dome in a 24-well plate

EBs were formed at a seeding density of 1000 cells/well in the ULA 384-well plate with the complete hESC medium as described above. From day 6, EBs were differentiated in the NIM in the ULA 384-well plate for an additional 6 days for NE formation. Each NE was mixed with 20 µL of 6 - 8 mg/mL Matrigel and then reseeded in a ULA 24-well plate to generate five of 20 µL Matrigel dome in each 24-well (single NE in each Matrigel dome). The Matrigel domes were incubated at 37°C for 20 minutes for complete gelation of Matrigel and then cultured with 400 µL of the NIM. The NIM was changed to the CDM-vit A after 2 days of culture, and the CDM-vit A was used for cerebral organoid differentiation until day 31 with medium change every alternate day.

### Assessment of EB and organoid viability

The viability of EBs and organoids were measured with a luminescence-based CellTiter-Glo 3D^®^ cell viability assay kit (G9681, Promega) by following the manufacturers’ recommended protocol. For the viability assessment of EBs in the ULA 384-well plate, after removing half the volume of the culture medium in the well and adding the same volume of the CellTiter-Glo^®^ reagent, the EBs were incubated with the 2-fold diluted reagent for 1 hour at room temperature on an orbital shaker. After 1 h incubation, 50 µL of the reagent solution was transferred to an opaque white 384-well plate, and luminescence intensity reading was obtained using Synergy H1 microplate reader (Agilent Technologies). Similarly, for the assessment of organoid viability on the pillar plate, the CellTiter-Glo reagent was diluted with the culture medium at 3:1 ratio and dispensed in the opaque white 384-well plate. The pillar plate with organoids was sandwiched onto the opaque white 384-well plate with the reagent, and the sandwiched plates were incubated at room temperature for 1 hour on the shaker. After 1 h incubation, the pillar plate was detached, and luminescence intensity reading was obtained from the opaque well plate using the microplate reader. The viability assessment with the CellTiter-Glo assay was performed three times to determine the reproducibility and uniformity of organoid generation on the 36PillarPlate.

In addition to the luminescence-based viability assay, a fluorescence-based cell viability assay was performed with calcein AM and ethidium homodimer-1. Briefly, organoids on the pillar plate on day 31 were rinsed with 1x PBS and stained with 2 μM calcein AM and 4 μM ethidium homodimer-1 in DMEM/F-12 medium for 2 hours at 37°C. After staining, the organoids were rinsed three times with PBS, and the fluorescence images were obtained with a fully automated bright-field and fluorescence microscope (BZ-X800E, Keyence).

### Immunofluorescence (IF) staining of whole cerebral organoids

All the solutions and reagents used for staining were dispensed in the 384DeepWellPlate at 80 µL/well. The pillar plate with cerebral organoids was sandwiched with the deep well plate to carry out organoid staining. Briefly, cerebral organoids on the pillar plate were rinsed with 1x PBS and treated with 4% paraformaldehyde (PFA) in the deep well plate overnight at 4°C. The organoids were permeabilized with 1% Triton X-100 in PBS for 1 hour at room temperature and then exposed to a blocking buffer (consisting of 4% normal donkey serum in 1x PBS with 0.5% Triton X-100) overnight at 4°C. Primary antibodies were diluted to recommended concentrations using the blocking buffer. The organoids were incubated with primary antibodies overnight at 4°C with gentle shaking. After primary antibody staining, organoids were rinsed with the blocking buffer three times and incubated with appropriate secondary antibodies diluted in the blocking buffer for 2 - 3 hours at room temperature or overnight at 4°C with gentle shaking. The stained organoids were washed with 0.5% Triton X-100 in PBS three times for 20 minutes each and incubated with 1 µg/mL DAPI in 0.5% Triton X-100 in PBS for 30 minutes. This was followed by final washing with PBS for 10 minutes three times and incubation with a tissue clearing solution (Visikol Histo-M) for 1 hour. Images were obtained with a confocal microscope (LSM710, Zeiss) with 10x magnification and were analyzed with ImageJ software. The primary and secondary antibodies used are listed in **Supplementary Tables 1 and 2**, respectively.

### Gene expression analysis *via* qPCR

The cerebral organoids in Matrigel were isolated from the pillar plate using Cultrex organoid harvesting solution (3700-100-01, R&D Systems) according to the manufacturer’s recommended protocol, which allows non-enzymatic depolymerization of Matrigel. Briefly, the pillar plate with cerebral organoids were sandwiched onto the deep well plate containing 80 µL of Cultrex organoid harvesting solution. The sandwiched plates were incubated for 30 minutes at 4°C and then centrifuged at 100 rcf for 10 minutes to detach the organoids. Total RNA was isolated from iPSCs and organoids on days 16 and 31 by using the RNeasy Plus Mini Kit (74134, Qiagen) with the manufacturer’s recommended protocol. Briefly, 10 - 12 organoids from each pillar plate were collected by using Cultrex organoid harvesting solution for qPCR analysis due to the small size of cerebral organoids on the pillar plate. cDNA was synthesized from 1 µg of RNA by following the protocol of cDNA conversion kit (4368814, Applied Biosystems). Real time PCR was performed using SYBR™ Green Master Mix (A25742, ThermoFisher) and forward/reverse primers from IDT Technology. The cycle was run 35 - 40 times at 95°C denaturation for 30 sec, 58 - 62°C annealing for 45 sec, depending on primer pair, and 72°C extension for 30 sec. The primers used are listed in **Supplementary Table 3**. We performed qPCR analysis three times to demonstrate reproducibility of organoid generation on the pillar plate.

### Statistical analysis

Statistical analysis was performed using GraphPad Prism 9.3.1 (GraphPad Software, Inc., CA, USA). The data was expressed as mean ± SD. P values were calculated using unpaired two-tailed t-test and one-way ANOVA. The statistical significance threshold was set at **** for p < 0.0001, *** for p < 0.001, ** for p < 0.01, * for p < 0.05, and ns = not significant (p > 0.05). Sample sizes are mentioned in the figure legends.

## Results

### Reproducibility of iPSC culture

Maintaining healthy and undifferentiated culture of iPSCs is crucial for the reproducible generation of brain organoids. EDi029A iPSC line was assessed for their uniformity in maintaining pluripotency and compact colonies across four different batches. We consistently observed the formation of compact colonies with smooth edges and higher nucleus to cytoplasmic ratio, throughout the culture period in multiple batches, demonstrating their applicability for uniform cerebral organoid generation **(Supplementary Fig. 1A)**. In addition, the iPSCs maintained their pluripotency with less than 10% differentiated cells when characterized for pluripotency with representative pluripotency makers such as OCT4 and SSEA4 (24,25) (**Supplementary Fig. 1B**).

### Improved viability of embryoid bodies (EBs) with CEPT cocktail

The differentiation of cerebral organoids can be initiated by forming EBs with Rho-kinase inhibitors such as CEPT and Y-27632 in the cell culture medium. These inhibitors enhance cell viability thereby potentially improving the consistency of organoid generation(11,26–28). Cellular stress during trypsinization and cryopreservation can hinder EB formation and compromise cell differentiation. In our study, we conducted a comparative analysis of the effects of recently developed CEPT and traditional Y-27632 on cell viability and uniform EB formation. The CEPT cocktail consisting of Chroman, Emricasan, Polyamine supplement, and Trans-ISRIB was developed to enhance cell viability after cryopreservation(26). Notably, the addition of CEPT in the culture medium led to the compact, round EBs with larger sizes, suggesting improved cell survival and proliferation compared to Y-27632 (**Fig. 1A & 1B**). Conversely, EBs formed with Y-27632 displayed more dead cell debris in their vicinity. The higher viability of EBs formed with CEPT was further confirmed through an ATP-based luminescence assay, as depicted in **Fig. 1C**. Based on these findings, we decided to use CEPT, rather than Y-27632, for EB formation in the cerebral organoid differentiation protocol.

**Figure 1.**
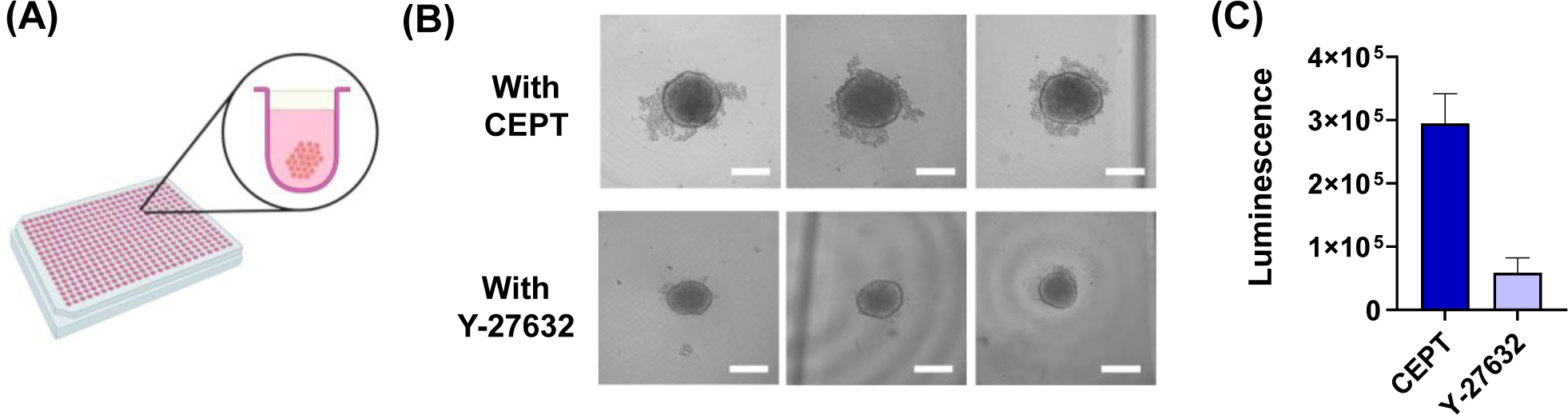
Formation of embryoid body (EB) in an ultralow attachment (ULA) 384-well plate: **(A)** Formation of single EBs with suspension of iPSCs in the ULA 384-well plate. **(B)** Morphology of day 7 EBs formed in hESC medium supplemented with CEPT or Y-27632 for 4 days. Scale bars: 200 μm. **(C)** Viability of EBs formed with CEPT or Y-27632 measured by using the ATP-based luminescence assay kit (n = 18).

### Robust transfer of EBs to the pillar plate *via* a sandwiching and inverting method

For brain organoid generation, individual neuroectoderm needs to be encapsulated in biomimetic hydrogels such as Matrigel and differentiated in a 6-/24-well plate, which is labor-intensive and error-prone. The ULA 384-well plate has been widely used for generating EBs, but it is challenging to use the ULA 384-well plate for organoid differentiation due to difficulty in changing growth media without disturbing cells in hydrogels and the necrotic core of organoids. To address this issue and improve the reproducibility of organoid formation, we developed a pillar plate platform to support culture of multiple spheroids/organoids simultaneously (**Fig. 2**). The method of spheroid transfer from the ULA 384-well plate to the pillar plate allowed us to efficiently load single spheroid on a one well-to-one pillar basis (**Fig. 3A**). The working principle of spheroid transfer is simple and straight-forward. The pillar plate with 5 µL of cold Matrigel can be sandwiched onto the ULA 384-well plate with EBs. The sandwiched plates can be immediately inverted to induce an EB from each well settled on each pillar. By incubating the sandwiched plates at 37°C in a CO_2_ incubator, EB transfer followed by Matrigel gelation can be achieved. The unique structure of the pillar plate with sidewalls and slits allowed robust spheroid transfer and encapsulation in Matrigel with minimal dilution. Cell culture media can be easily changed by adding fresh media in a deep well plate and sandwiching the pillar plate with EBs encapsulated in Matrigel onto the deep well plate. In addition, *in situ* imaging of spheroids and organoids on the pillar plate can be performed due to the small dimension and the fixed location of the encapsulated cells.

**Figure 2.**
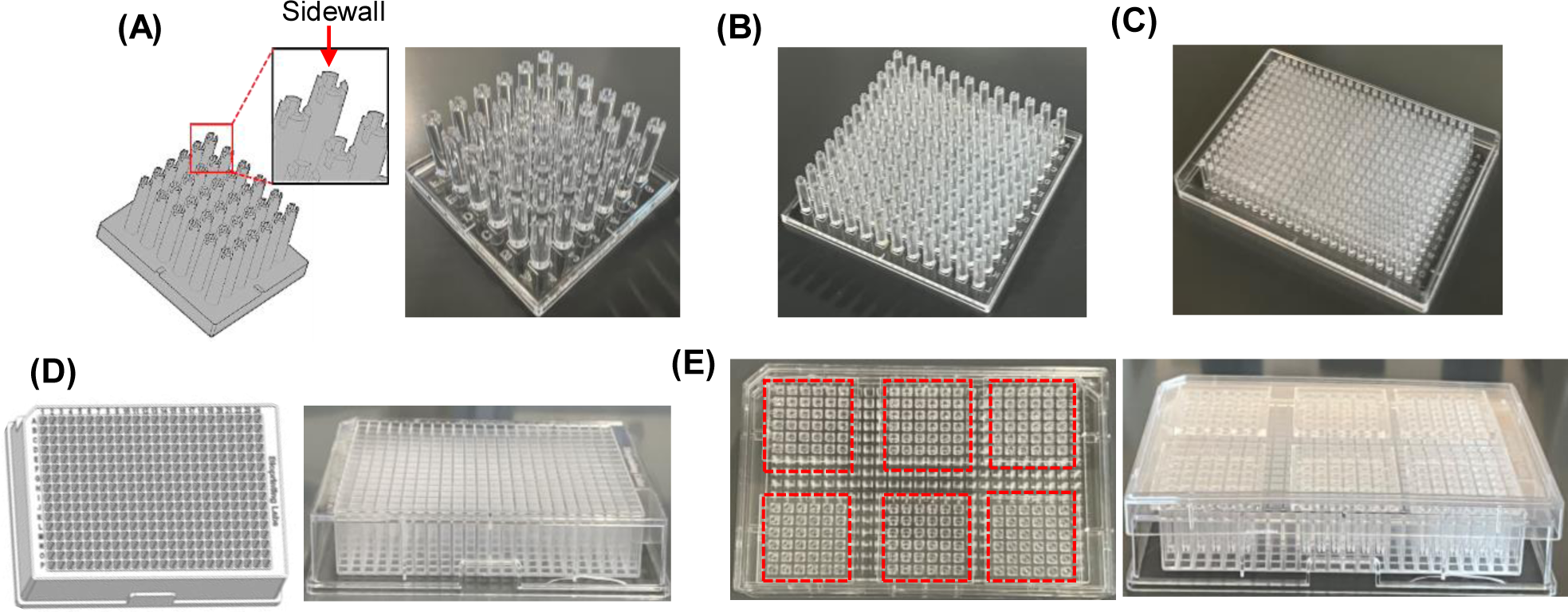
Injection-molded pillar plates and the complementary deep well plate for static organoid culture: **(A)** SolidWorks design (left) and picture of the injection-molded 36PillarPlate with 6 x 6 array of pillars (right). **(B)** The injection-molded 144PillarPlate with 12 x 12 array of pillars. **(C)** The injection-molded 384PillarPlate with 16 x 24 array of pillars. **(D)** SolidWorks design (left) and picture of the injection-molded 384DeepWellPlate with 16 x 24 array of deep wells (right). (D) Six of the 36PillarPlate sandwiched onto the 384DeepWellPlate for static organoid culture: top view (left) and side view (right).

**Figure 3.**
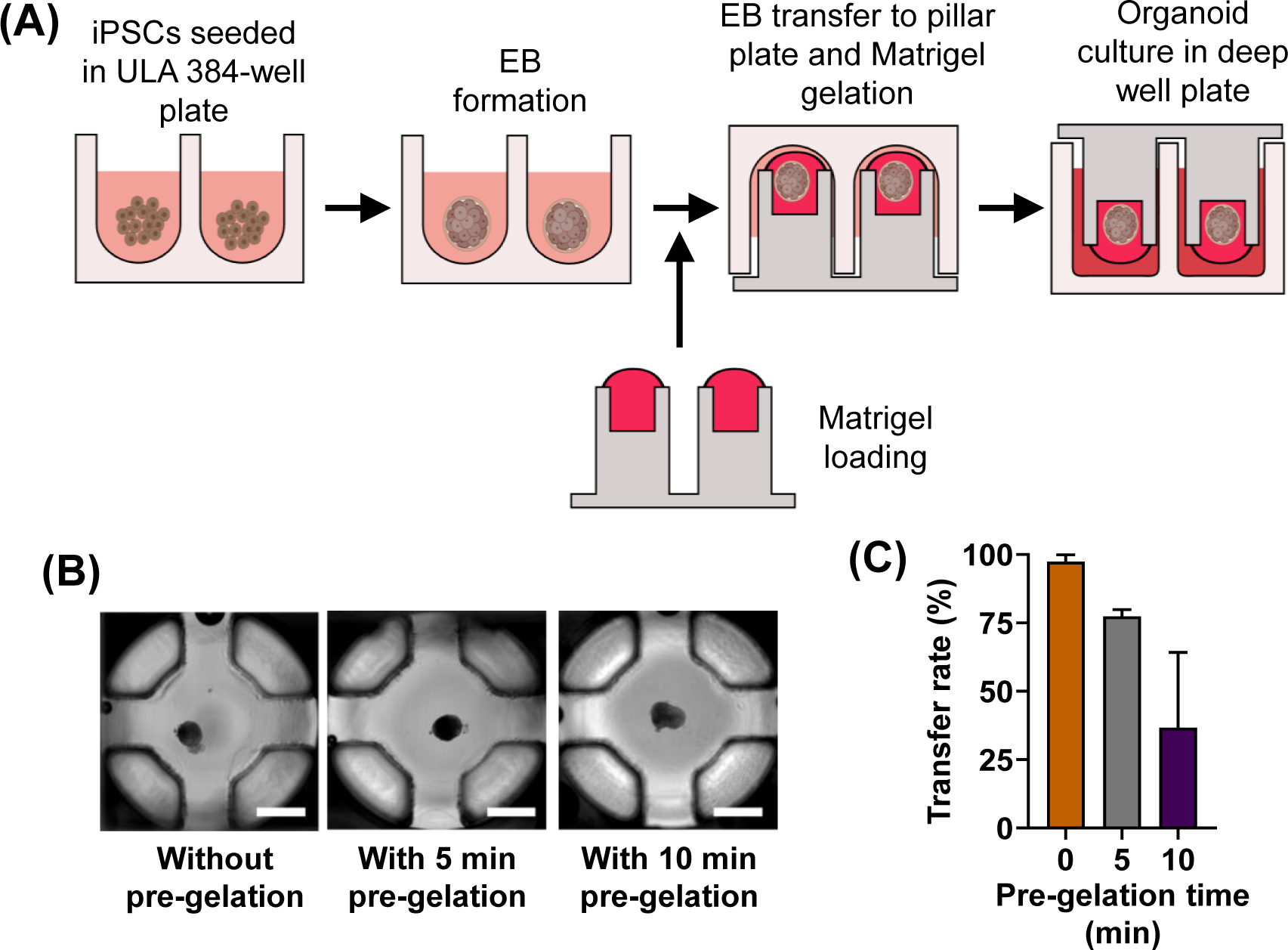
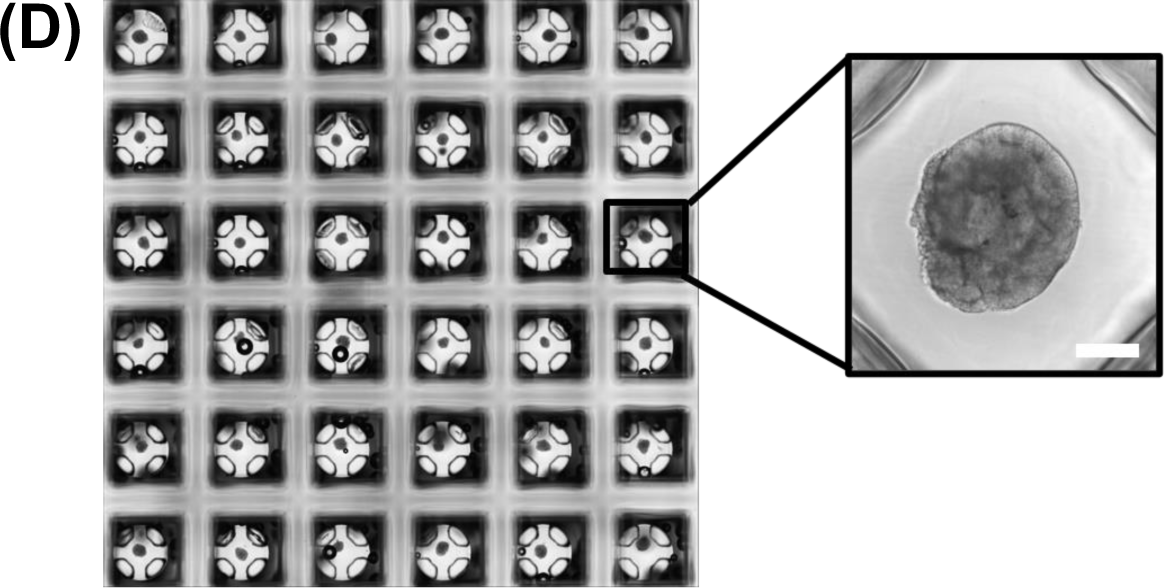
Optimization of spheroid transfer to the 36PillarPlate from the ULA 384-well plate: **(A)** Steps necessary to transfer single spheroids from the ULA plate to the pillar plate. **(B)** Representative images of iPSC spheroids transferred at different pre-gelation time of Matrigel. Scale bars: 500 μm. **(C)** The success rate of iPSC spheroid transfer at different pre-gelation time. n = 18. **(D)** Stitched images of EBs encapsulated in Matrigel on the pillar plate after transfer. Scale bar: 200 µm.

For optimization of the spheroid transfer protocol, several parameters including incubation time at 37°C for Matrigel gelation, spheroid size, and pre-gelation of Matrigel at room temperature were tested to achieve high success rates of spheroid transfer. Among the incubation time at 37°C tested, 30 – 40 minute incubation of the sandwiched plates was necessary to have complete Matrigel gelation on the pillar plate in the ULA 384-well plate. The effect of the spheroid size on transfer efficiency was tested with ReNcell spheroids with a cell seeding density in a range of 500 - 9000 cells/well in the ULA 384-well plate. The size of spheroids formed was between 120 - 350 μm after 4 days of culture with the seeding density (**Supplementary Fig. 2A**). The success rate of spheroid transfer was > 95% at all the size of spheroids tested on the pillar plate with 4 - 6 mg/mL of Matrigel and 20 - 30 minutes of Matrigel gelation at 37°C in the CO_2_ incubator (**Supplementary Fig. 2B**). This result indicates that the speed of spheroid precipitation on the pillar plate is much faster than that of Matrigel gelation, leading to successful encapsulation of spheroids. Since cerebral organoids differentiated from EBs at 1000 cells/well seeding density showed more complex morphology as compared to those generated with 3000 cells/well (**Supplementary Fig. 3**), the established spheroid transfer protocol can be used for cerebral organoid culture. The Matrigel concentration for EB encapsulation was increased to 6 - 8 mg/mL to account for potential dilution of Matrigel after sandwiching the pillar plate into the ULA 384-well plate during spheroid transfer. Finally, partial gelation of Matrigel on the pillar plate at room temperature (i.e., pre-gelation) before spheroid transfer was investigated due to the concern for potential Matrigel dilution in the medium in the ULA 384-well plate during the sandwiching (**Fig. 3B & 3C**). A high success rate of EB transfer was obtained when the pillar plate with Matrigel was sandwiched into the ULA 384-well plate without pre-gelation, indicating that premature polymerization of Matrigel prohibits successful spheroid transfer and encapsulation on the pillar plate (**Fig. 3D**).

### Generation of cerebral organoids on the pillar plate

Cerebral organoids were generated on the pillar/deep well plate by following the Lancaster protocol (29) with slight modification for early embedding of EBs in Matrigel on the pillar plate (**Fig. 4A**). We selected the Lancaster protocol because it has been validated by numerous researchers and is well established for cerebral organoid culture as compared to other protocols developed recently. The differentiation process of cerebral organoids started with EB formation and transfer on the pillar plate, followed by neural induction, neural expansion, and maturation without the use of any exogenous factors. EBs formed in the ULA 384-well plate showed clear borders that were visible on day 2 and were brighter on day 4 (**Fig. 4B**). EBs showed brighter morphology with smooth edges on the day of transfer on day 7. iPSCs can be stimulated to develop germ layers including ectoderm within the aggregates known as EBs and can be developed into cerebral organoids with low bFGF in hESC medium (11). After transferring on the pillar, they were cultured in the neural induction medium for 5 - 6 days which led to uniform neuroectoderm (NE) formation by the suppression of mesodermal tissue development(11). The organoids were then cultured in cerebral organoid differentiation medium (CDM) without vitamin A until day 31. Cerebral organoids at early stages (day 15 - 20) on the pillar plate showed several small neural rosettes, which are reminiscent of neuroepithelia (**Fig. 4B**), similar to previously published data (11). CDM without vitamin A supports the developmental progression of neural progenitors and their progeny (11). Neuronal development and differentiation are supported by neurobasal medium and B27 supplement (11). Additional supplements such as 2-mercaptoethanol and insulin maintained the neural stem cells (NSCs)(30). They were cultured until day 31 only, which was enough to observe some of the representative markers related to cerebral organoids for analyzing their formation on the pillar plate. These cerebral organoids generated on the pillar plate have expression levels of crucial genes related to brain development, similar to that from conventional Matrigel dome culture (**Supplementary Fig. 4**) with slight increment in most of the genes, demonstrating the compatibility of the pillar/deep well plate for robust cerebral organoid culture (**Supplementary Fig. 5**).

**Figure 4.**
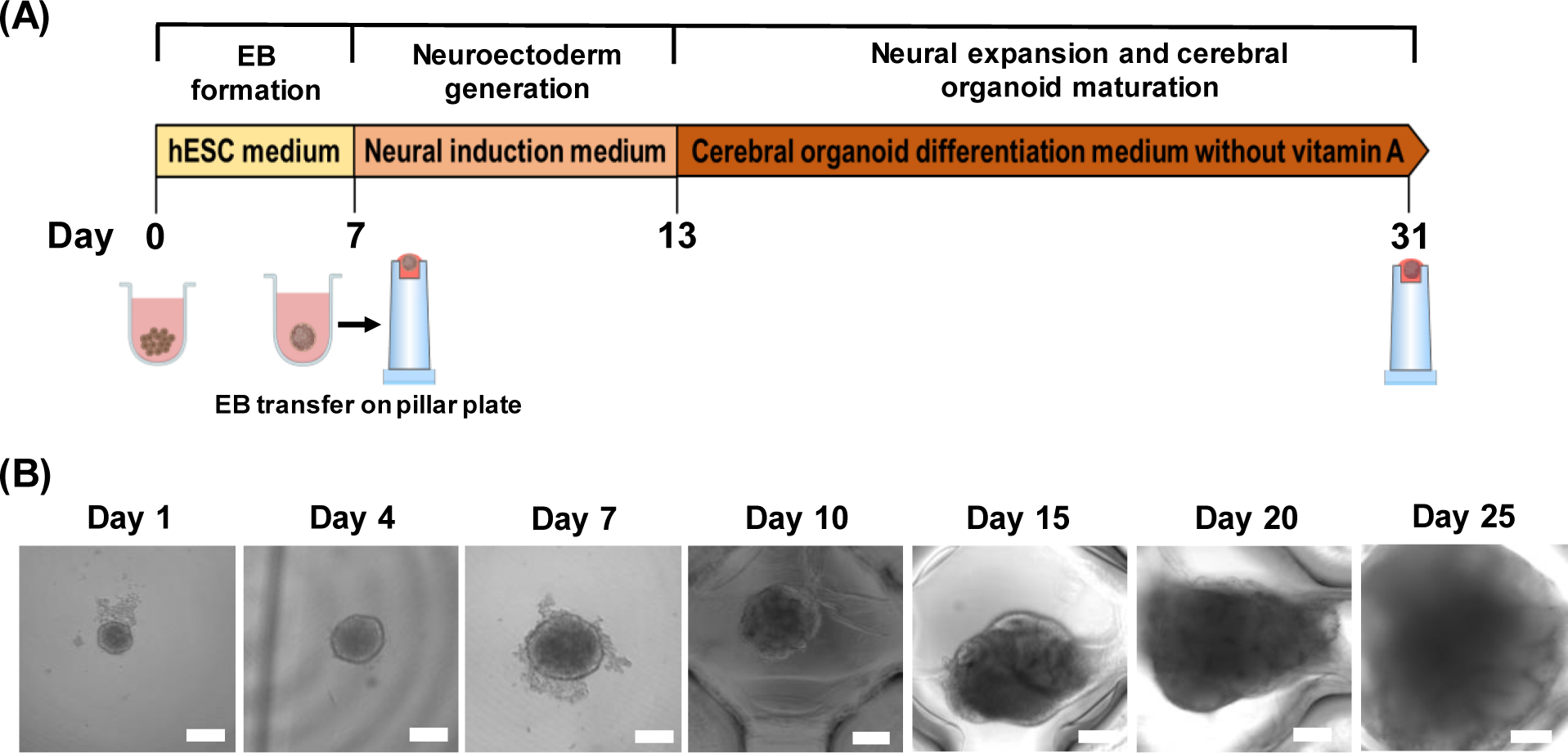
Generation of cerebral organoids on the pillar plate using EDi029A iPSCs: **(A)** Differentiation process of cerebral organoids on the pillar plate. **(B)** Cerebral organoids generated on the pillar plate over time. Scale bars: 200 μm.

### Characterization of cerebral organoids generated on the pillar plate

The cerebral organoids generated on the pillar plate were characterized by qPCR and immunofluorescence staining. Gene expression analysis through qPCR on day 16 and 31 of differentiation revealed significant changes in marker expression over time, as illustrated in **Figure 5**. By day 16 of differentiation, the organoids exhibited a significant reduction of over 1000-fold in pluripotency marker *OCT4*, indicating the initiation of differentiation. Simultaneously, there was a notable, more than 30-fold increase in the expression of the neural progenitor marker, paired box protein 6 (*PAX6*), signifying successful neural induction. These changes align with previously published data(10). *PAX6* expression was significantly higher throughout the differentiation up to day 31. The forebrain marker, forkhead box protein G1 (*FOXG1*) displayed a 10-fold increase by day 16, maintaining their elevation until day 31, reflecting earlier brain regionalization. Similarly, the neuronal marker, microtubule-associated protein 2 (*MAP2*) exhibited a significant increase from nearly 20-fold to 50-fold between day 16 and day 31, indicating the maturation of brain organoids over time. The expression of the cortical plate precursor and intermediate progenitor marker, T-box protein (*TBR2*) decreased from 30-fold to 20-fold from day 16 to day 31, possibly due to the transition of intermediate progenitor cells into post mitotic neurons (31). The neural stem cell marker, *SOX2*, was consistently expressed at a 2.5-fold level on both day 16 and day 31. Deep cortical neuronal marker, COUP-TF-interacting protein 2 (*CTIP2*) and choroid plexus marker, *TTR* were expressed by 3.5-fold only on day 31 following a pattern observed in previous study (11). The neuronal marker *TUBB3* was expressed at 10-fold on day 16 and increased to 16-fold on day 31. *TBR1*, a preplate marker, was expressed at 115-fold on day 16, and then increased to 330-fold on day 31, indicating the formation of the subventricular zone. Overall, the cerebral organoids generated on the pillar plate recapitulated key aspects of neurogenesis, cortical organization, and brain regionalization, revealing early features of brain development.

**Figure 5.**
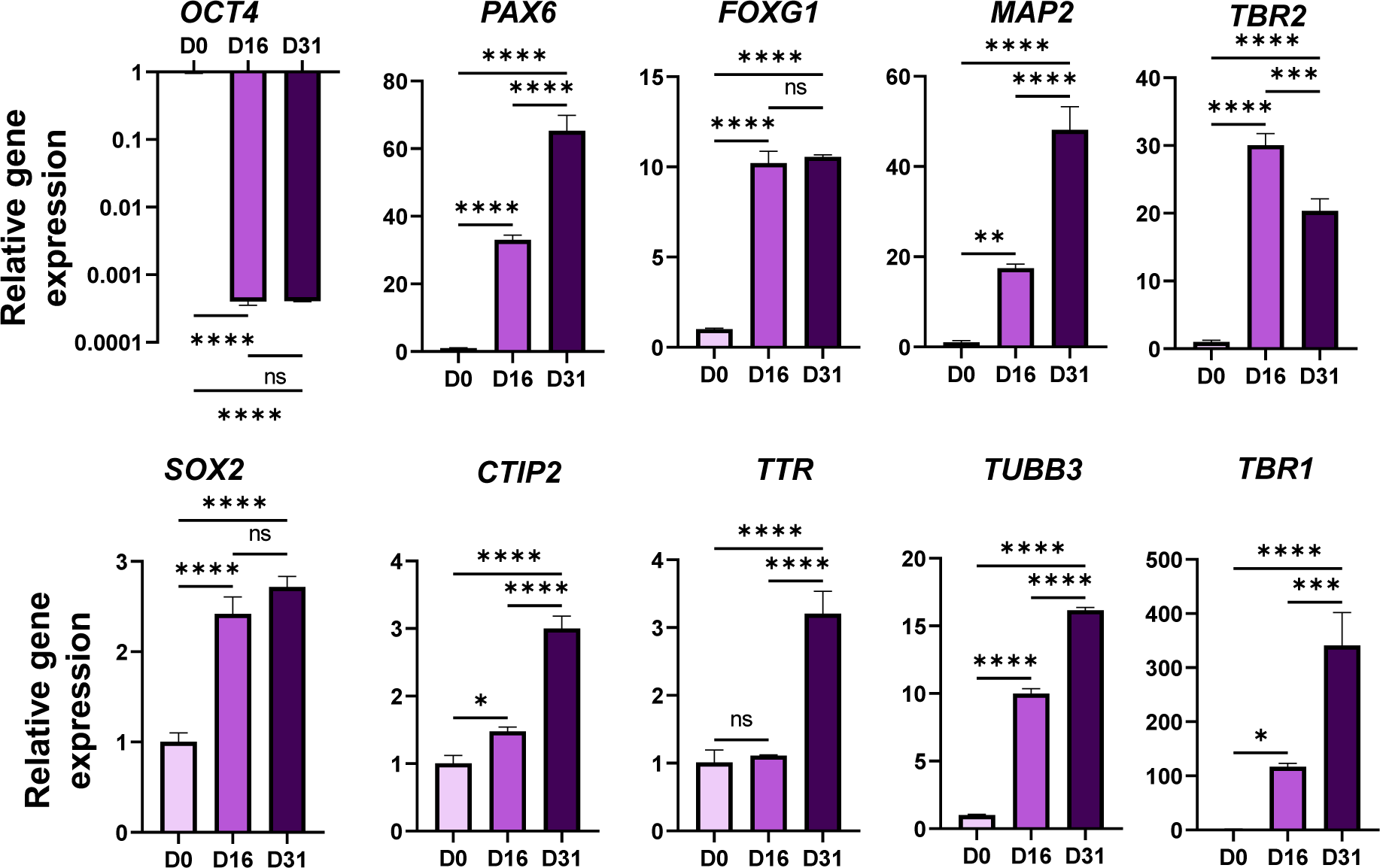
Characterization of days 16 and 31 cerebral organoids on the pillar plate. Gene expression of *OCT4* pluripotency marker, *PAX6* forebrain neuroprogenitor marker, *FOXG1* forebrain marker, *MAP2* mature neuronal marker, *TBR2* intermediate progenitor marker, *SOX2* proliferating neuroprogenitor marker, *CTIP2* deep cortical neuronal marker, *TTR* choroid plexus marker, *TUBB3* neuronal cytoplasm marker, and *TBR1* preplate marker analyzed by qPCR. Statistical significance was performed by one-way ANOVA. **** for p < 0.0001, *** for p < 0.001, ** for p < 0.01, * for p < 0.05, and ns = not significant (p > 0.05). n = 10 - 12 per qPCR run.

In addition to qPCR analysis, immunofluorescence staining was performed on day 31 to determine whether the cells were randomly scattered or developed with discrete regions within whole cerebral organoids (**Fig. 6A**). The stained organoids on day 31 revealed the high expression level of several forebrain region biomarkers, including *PAX6*, *MAP2*, *SOX2*, and *CTIP2* (10). To demonstrate the initiation of cortical organization, we stained the organoids with SOX2 and CTIP2. The distinct separation of SOX2+ and CTIP2+ cells could represent the presence of the ventricular zone and the cortical plate-like region as defined in the previous study (32). SOX2+ cells are present in the ventricular zone whereas CTIP2+ cells are in the deep cortical plate. The presence of these markers was distinct in the organoids (**Fig. 6A**), which could support the initiation of cortical plate layer formation within the brain organoids. Since proper cortical layering would not be initiated in 31 days of differentiation, we did not stain the organoids with SATB2, which is expressed in the late stage of differentiation. In addition to the localized expression of FOXG1 forebrain region marker in the organoids, the organoids were also stained with TBR2 intermediate progenitor marker, which is also a precursor of cortical plate (11). A few TBR2+ cells were dispersed at the periphery of the ventricular zone. MAP2 mature neuronal marker revealed the appearance of the basal neural layer that represents a superficial preplate zone-like layer in day 31 organoids (**Fig. 6A**). The quantitative expression of fluorescence intensity obtained from different markers within the cerebral organoids on day 31 was provided in **supplementary Fig. 6**.

**Figure 6.**
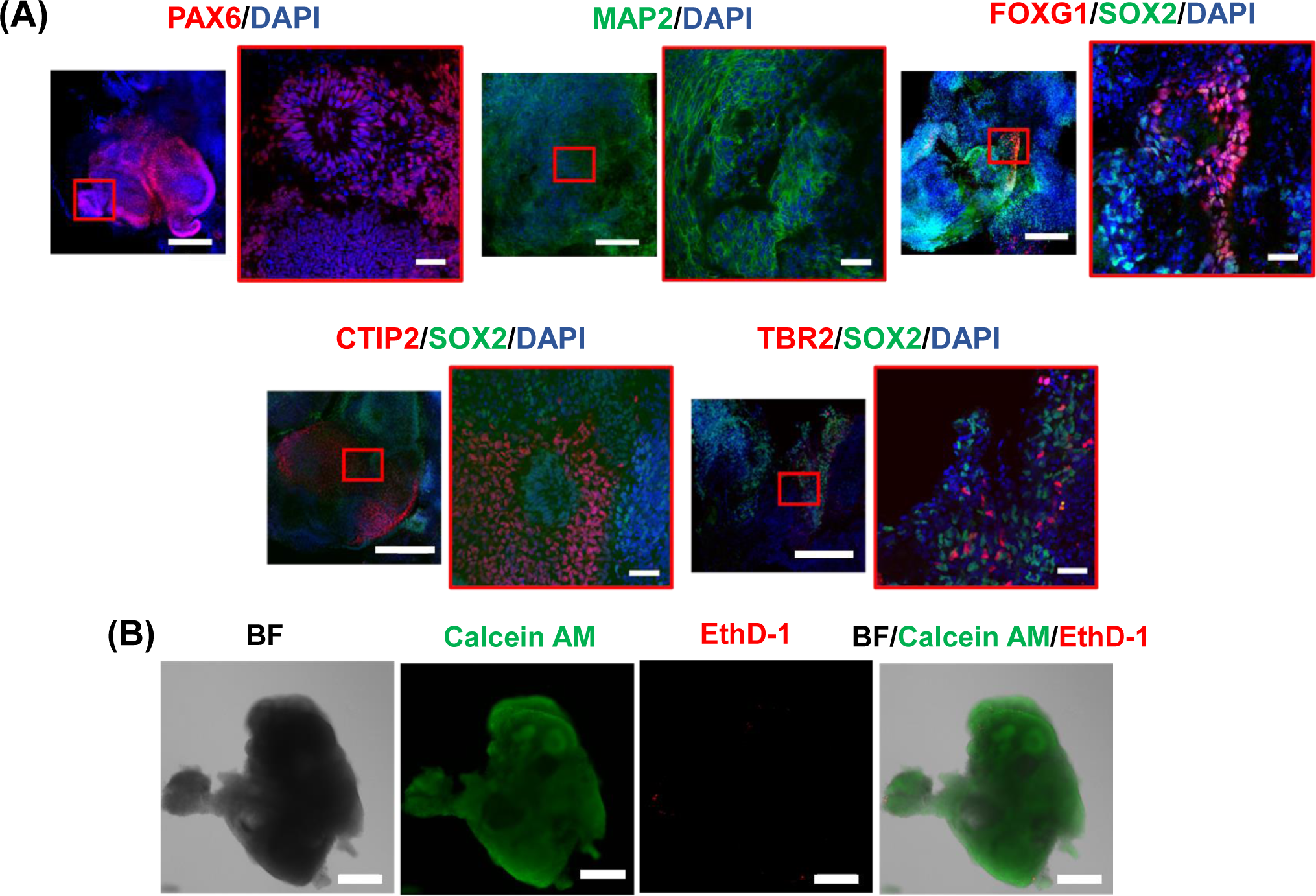
Characterization of day 31 cerebral organoids on the pillar plate: **(A)** Immunofluorescence staining of cerebral organoids for PAX6 forebrain neuroprogenitor marker, MAP2 mature neuronal marker, FOXG1 forebrain marker, SOX2 proliferating neuroprogenitor marker, CTIP2 deep cortical neuronal marker, and TBR2 intermediate progenitor marker. Scale bars: 200 µm and 50 µm (magnified). **(B)** The viability of cerebral organoids assessed with calcein AM and ethidium homodimer-1 (EthD-1) staining. Scale bars: 200 µm.

Moreover, live/dead cell staining of day 31 cerebral organoids on the pillar plate with calcein AM and ethidium homodimer-1 (EthD-1) showed very few dead cells (**Fig. 6B**). This result indicates that the pillar/deep well plate platform can effectively support the culture of cerebral organoids by allowing the sufficient diffusion of nutrients and oxygen to the core due to the small size.

### Reproducibility of cerebral organoid generation on the pillar plate

One of the significant technical challenges in current cerebral organoid protocols is the lack of consistency and uniformity, which presents a major obstacle for high-throughput assessment of compounds with organoids. In this study, we overcome this limitation by consistently generating uniform cerebral organoids on the pillar plate, using a method that involved transferring single EB to each pillar. The EBs formed uniformly in the ULA 384-well plate (**Fig. 7A**) were simultaneously transferred to the pillar plate by the simple sandwiching and inverting method (**Fig. 3D**). Due to single, uniform size of EB transferred to the pillar plate, we were able to achieve uniform and reproducible cerebral organoid generation with the coefficient of variation (CV) values between 7 - 19% in three replicates (**Fig. 7B & 7D**), which was measured by the ATP-based cell viability assay. In every batch of culture, the organoids showed expanded neuroepithelial bud outgrowth (thin arrow) that is optically clear and surrounded by a visible lumen which is the typical morphology of cerebral organoids (**Fig. 7C**), indicating the successful differentiation as seen in previous study (11,21). Similarly, the gene expression level of cerebral organoids analyzed from three different plates showed similar expression levels on day 16 of differentiation (**Fig. 7E**).

**Figure 7.**
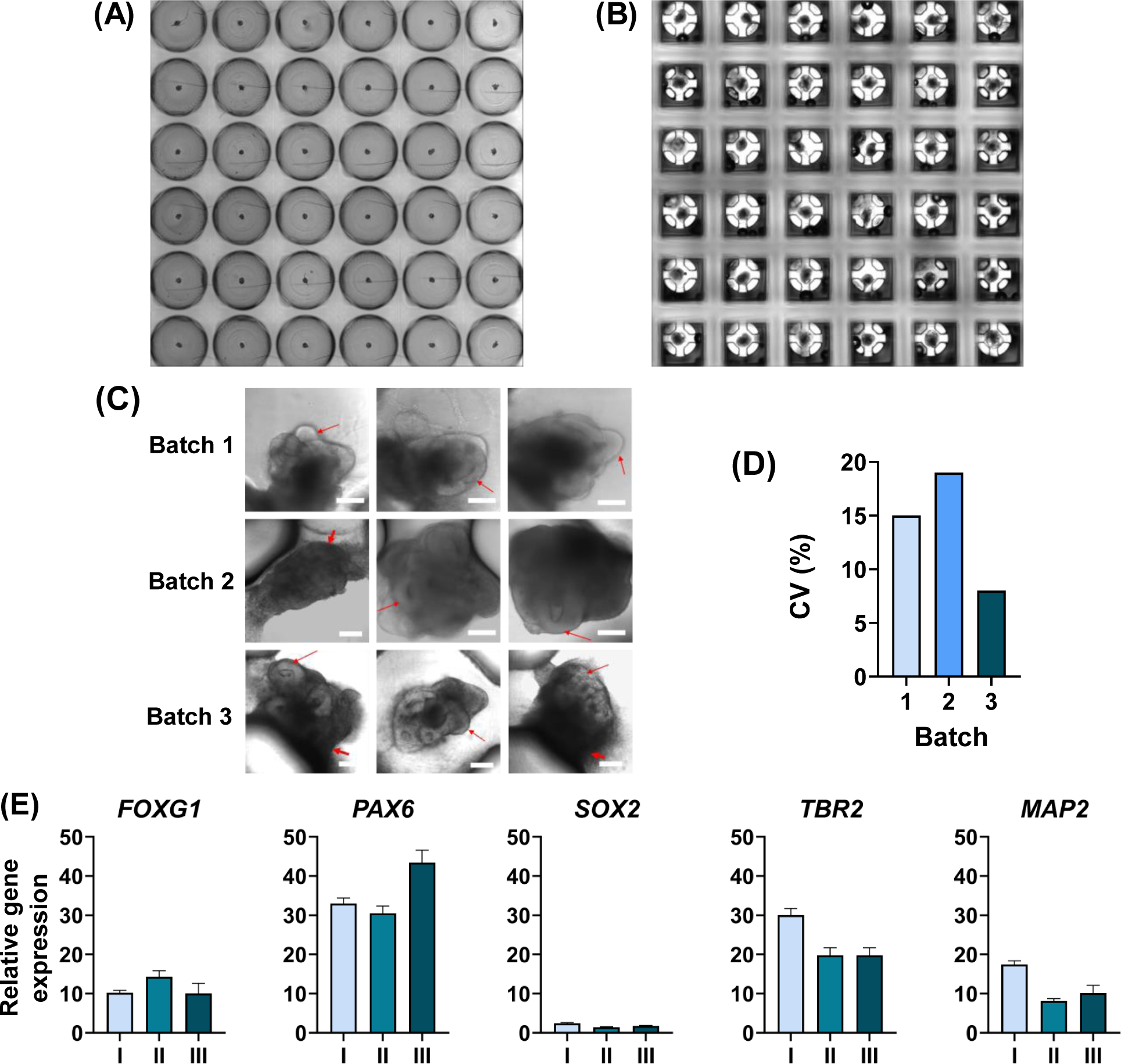
Reproducible generation of cerebral organoids on the pillar plate: **(A)** Stitched images of EBs in the ULA 384-well plate. **(B)** Stitched images of day 31 cerebral organoids on the pillar plate. **(C)** Reproducibility of early cerebral organoids with expanded neuroepithelium on the pillar plate from triplicate experiments. Early cerebral organoids generated on the 36PillarPlate showed expanded neuroepithelial bud outgrowth (thin arrow) that is optically clear and surrounded by a visible lumen. Other outgrowing and migrating cells are also visible (thick arrow) that are not neuroepithelial. Magnification: 10x, Scale bars: 200 µm. **(D)** The CV value of organoid viability measured with the ATP-based luminescence assay. n = 18. The CV values were in the range of 7 - 19% from triplicate trials, indicating the robustness of cerebral organoid generation on the pillar plate. **(E)** Changes in gene expression levels in day 16 cerebral organoids obtained from triplicate trials.

## Discussion

Although brain organoids have the potential to replicate the architecture and function of the brain, their development and application are significantly hampered by issues related to variability, technical complexity, limited scalability, and high costs (1). Currently, brain organoids are typically cultured in 6-/24-well plates, petri dishes, and spinner flasks, which require a large volume of cell culture media, growth factors, and additives (1,15,18). In addition, the culture process involves labor-intensive and cumbersome steps of encapsulation and dissociation of spheroids and organoids in hydrogels, resulting in low throughput along with a higher risk of contamination. For instance, in studies conducted by Lancaster et al.(11), Zhang et al.(33), and Ham et al.(34), EBs formed in ULA well plates were manually transferred one by one onto dimples of parafilm using a 200 µL pipette. Subsequently, a droplet of Matrigel was added to each one of EBs, and then they were cultured either in a petri dish by placing on an orbital shaker, or in a spinning bioreactor containing 75 - 100 mL of cell culture media. In the study by Bodnar et al.(1), organoids were collected in a conical tube, resuspended in Matrigel, and cultured in a ULA 6-well plate. Similarly, Qian et al.(32) generated region-specific brain organoids using a miniature spinning bioreactor, designed for small-scale organoid generation to enhance the diffusion of nutrients and oxygen into the core of the organoid. However, this method requires a laborious setup of the miniature bioreactor, even though it could reduce the use of cell culture media. In addition, microfluidic devices have been incorporated into brain organoid culture to provide a dynamic environment and to enhance the organoid maturity. Nevertheless, they also involve cumbersome steps of transferring individual spheroids and organoids encapsulated in biomimetic hydrogels into each organoid culture chamber (19,35). Furthermore, microfluidic devices are inherently low throughput due to the need of pumps and tubes rendering them user-unfriendly. In an effort to generate consistent cerebral organoids, Sivitilli et al.(36) implemented a “V” bottom, nonadherent, 96-well plate instead of a surface-treated, “U” bottom, polystyrene plate for robust EB formation. However, the workflow in this study also involved manual embedding of EBs in Matrigel, followed by culturing them in an orbital shaker. In summary, miniature organoid culture is highly beneficial for saving cell culture media, performing *in situ* organoid imaging, and reducing the necrotic core of organoids. However, it is challenging to differentiate organoids in high-density well plates such as a 384-well plate due to difficulty in changing growth media without disturbing cells in hydrogels and the necrotic core of organoids.

In this study, we addressed the technical challenges in brain organoid generation, which include the cost of cell culture media, reproducibility and scalability of organoid culture and analysis, and complexity of the culture process. Using our spheroid transfer and encapsulation method on the pillar plate, we simplified the EB encapsulation in hydrogel and organoid culture process with high reproducibility. EBs formed in the ULA 384-well plate can be transferred to the pillar plate on one EB-to-one pillar basis by simple sandwiching and inverting with minimal manual intervention. Changing cell culture media during differentiation is simple and straightforward without any concerns for disrupting EBs in hydrogels. Fresh media (80 µL/deep well) can be dispensed into the 384DeepWellPlate using a multichannel pipette or a liquid dispenser, and then the pillar plate with organoids in Matrigel can be sandwiched onto the 384DeepWellPlate all at once for media change. Since the pillar plate requires EBs in 5 µL of Matrigel spot and 80 µL of cell culture media in the 384DeepWellPlate, the cell culture volume necessary for organoid culture is 10 - 100-fold reduced, as compared to conventional 6-/24-well plates, petri dishes, and spinner flasks. The reduced cell/Matrigel volume could enhance the diffusion of nutrients and oxygen to the core of organoids, potentially leading to high maturity of organoids as compared to conventional Matrigel dome culture (**Supplementary Fig. 5**). The maturity of organoids could be further enhanced by incorporating dynamic culture in the pillar/perfusion plate (37,38). In addition, organoid staining and analysis can be streamlined on the pillar plate using existing lab equipment. For example, immunofluorescence staining of cerebral organoids with antibodies as well as cell viability assays with fluorescent dyes were performed simultaneously with organoids on the pillar plate and reagents in the deep well plate (**Fig. 6**). Due to the small scale, organoid analysis can be done in high throughput with existing fluorescence microscopes and microtiter well plate readers for performing high-throughput organoid imaging and acquiring absorbance, fluorescence, and luminescence signals from organoids. The throughput of organoid-based assays can be further enhanced by using the 144PillarPlate with 144 pillars or the 384PillarPlate with 384 pillars per plate, instead of using the 36PillarPlate with 36 pillars. Furthermore, the reproducibility of brain organoid generation was demonstrated with the CV values below 20% by using the spheroid transfer and encapsulation method on the pillar plate (**Fig. 7B**). Therefore, the pillar/deep well plate platform could reduce the cost of cell culture media, enhance reproducibility and scalability of organoid culture and analysis, and simplify the organoid culture process with minimal manual intervention.

There is room for improvement of the spheroid transfer and encapsulation method on the pillar plate. Currently, we load 6 - 8 mg/mL of diluted Matrigel on the pillar plate with a multichannel pipette, which takes approximately 2 minutes for the 36PillarPlate. Aligning pipette tips to the pillars is quite cumbersome and often leads to premature gelation of Matrigel and failure of spheroid transfer, particularly when undiluted Matrigel is used. To avoid this issue, a cell loading plate is under development (**Supplementary Fig. 7**), which allows hydrogels alone or cells suspended in hydrogels loaded on the pillar plate by simple one-step stamping. The unique structure of the pillar with sidewalls and slits allows hydrogels alone or cells in hydrogels loaded reproducibly by stamping. The second issue pertains to the potential detachment of organoid spots from the pillar plate during long-term cell culture which might be due to surface incompatibility between alginate coating and Matrigel on the pillar. Alginate was used to prevent 2D cell growth on the surface of the pillars during long-term cell culture due to its low affinity to the integrin-binding domain on the cell surface. With current alginate coating, up to 5 - 10% of EBs in Matrigel could be detached from the pillar plate within a few days of culture, and the remaining 90 - 95% of cell spots were attached even after 31 days of culture and Matrigel remodeling by organoids (**Fig. 7**). This problem could be resolved by adding alginate in Matrigel or using other non-adherent polymer coatings that have more affinity to Matrigel. Dilution of Matrigel on the pillar plate in the ULA 384-well plate during spheroid transfer and encapsulation would not be a concern because of the pillar located very close to the bottom of the ULA well with small volume of media available and relative fast gelation of small volume of Matrigel (5 µL). The last remaining issues are related to immaturity of brain organoids due to relatively short-term culture and the potential necrotic core of organoids due to diffusion limitation of nutrients and oxygen during long-term cell culture. Brain organoids could not be fully mature in 31 days of differentiation. Thus, proper cortical layering and high expression of late-stage biomarkers such as SATB2 could not be achieved in 31 days of differentiation. It would be necessary to differentiate brain organoids for 2 – 4 months to observe relatively high maturity. For long-term culture, brain organoids have been transferred to dynamic culture conditions in petri dishes on an orbital shaker or in spinner flasks after a few weeks to one month of initial culture in Matrigel domes due to the necrotic core issue. The pillar plate can be coupled with a perfusion well plate for long-term, dynamic organoid culture while reducing the concern for the necrotic core (37,38).

## Conclusions

In summary, we developed a simple and robust method of spheroid transfer and encapsulation in hydrogel on the pillar plate, which could reduce the cost of cerebral organoid culture by 10 - 100-fold miniaturization, enhance reproducibility and scalability of organoid culture and analysis, and simplify the organoid culture process with minimal manual intervention. The spheroid transfer and encapsulation method on the pillar plate can be used for general static 3D cell culture including spheroids and organoids. Conventional methods of spheroid and organoid transfer and encapsulation are tedious, labor-intensive, and thus low throughput. This step is often the critical bottleneck in scale-up organoid culture and analysis for high-throughput screening of compounds with organoids. We envision that our spheroid transfer and encapsulation approach on the pillar plate could be widely adopted in academic and industry settings due to its simplicity and applicability to any 3D cell culture.

## Acknowledgement

This study was supported by the National Institutes of Health (NCATS R44TR003491 and NIDDK UH3DK119982).

## Supplementary Information

**Supplementary Table 1.** List of primary antibodies.

| Name | Species | Dilution | Vendor | Catalog # |
| --- | --- | --- | --- | --- |
| PAX6 | Rabbit | 1:350 | Abcam | ab195045 |
| FOXG1 | Rabbit | 1:100 | Abcam | 196868 |
| SOX2 | Mouse | 1:40 | Santa Cruz<br>Biotech | sc-365823 |
| TBR2 | Rabbit | 1:200 | Abcam | 183991 |
| CTIP2 | Rabbit | 1:200 | Abcam | ab 240636 |
| MAP2 | Rabbit | 1:200 | Abcam | 183830 |

**Supplementary Table 2.** List of secondary antibodies.

| Name | Dilution | Vendor | Catalog # |
| --- | --- | --- | --- |
| Donkey anti-Rabbit IgG (H+L) Highly Cross-Adsorbed Secondary Antibody, Alexa Fluor 647 | 1:200 | ThermoFisher | A-31573 |
| Donkey anti-Mouse IgG (H+L) Highly Cross-Adsorbed Secondary Antibody, Alexa Fluor 488 | 1:200 | ThermoFisher | A-21202 |
| Donkey anti-Rabbit IgG (H+L) Highly Cross-Adsorbed Secondary Antibody, Alexa Fluor 488 | 1:200 | ThermoFisher | A21206 |

**Supplementary Table 3.**
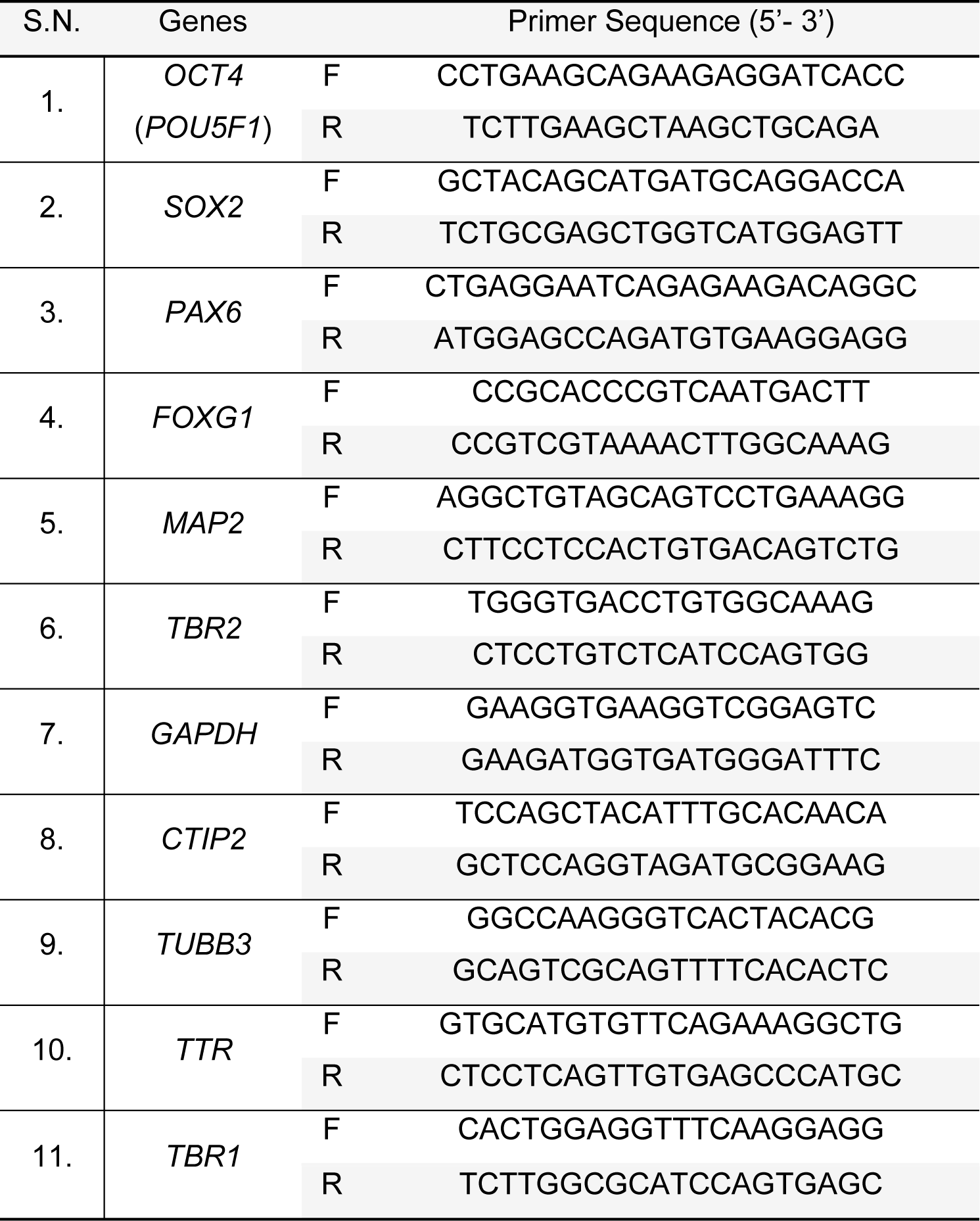
List of primers.

**Supplementary Figure 1.**
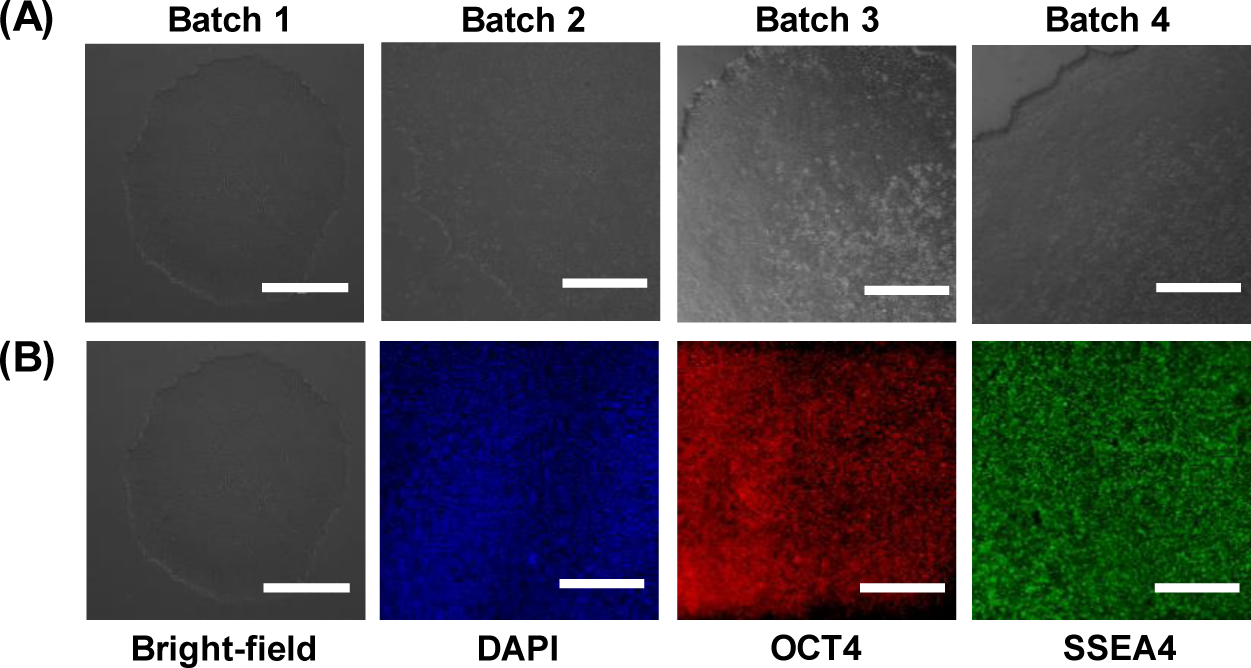
Culture and characterization of EDi029A iPSC line: **(A)** Brightfield images of iPSC colony from different batches. **(B**) Immunofluorescence images of iPSC colony showing the expression of pluripotency markers, including OCT4 and SSEA4. Magnification: 10x, Scale bar: 400 µm.

**Supplementary Figure 2.**
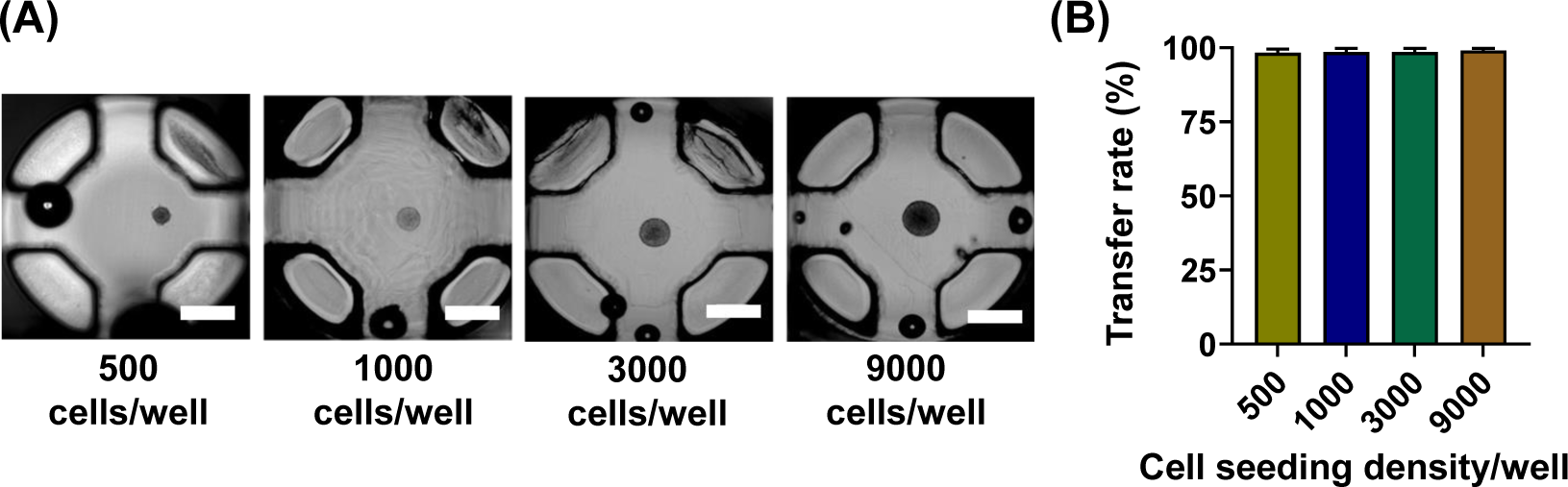
Optimization of spheroid transfer to the 36PillarPlate from the ULA 384-well plate: **(A)** Brightfield images of ReNcell spheroids with different sizes transferred to the pillar plate. Scale bar: 500 μm. **(B)** The success rate of ReNcell spheroid transfer at different seeding density. n = 18.

**Supplementary Figure 3.**
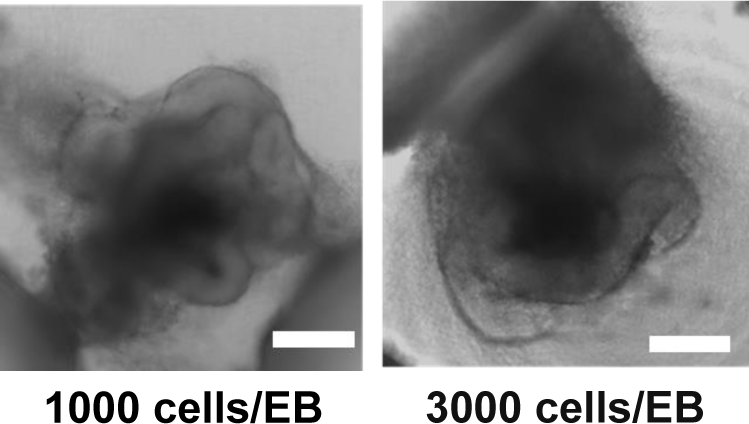
Brightfield image of day 16 cerebral organoids on the pillar plate with a seeding density of 1,000 (left) and 3,000 (right) cells per EB. Scale bars: 200 μm.

**Supplementary Figure 4.**
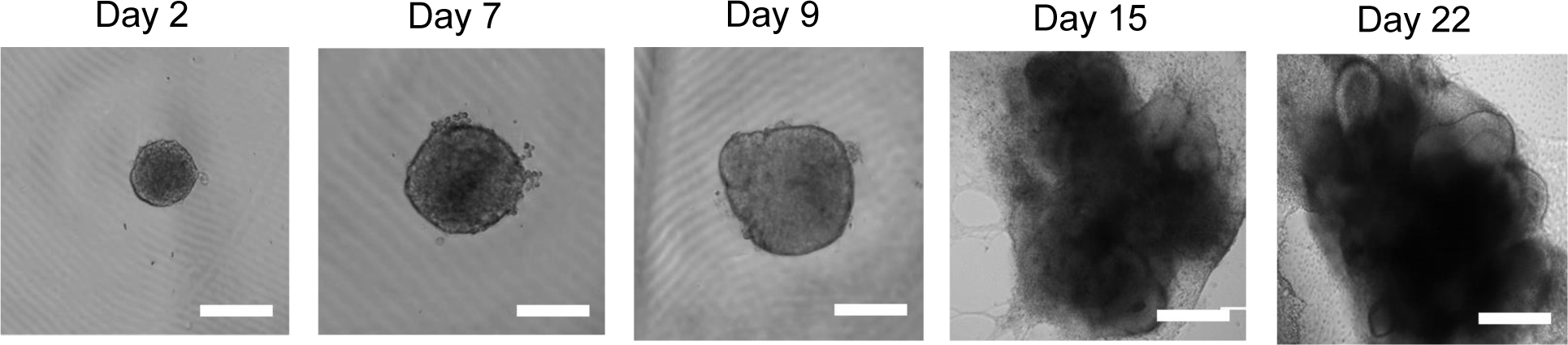
Generation of cerebral organoids in Matrigel domes in a 24-well plate. Scale bars: 200 µm. Two-dimensional cell growth was unavoidable in Matrigel domes in the 24-well plate.

**Supplementary Figure 5.**
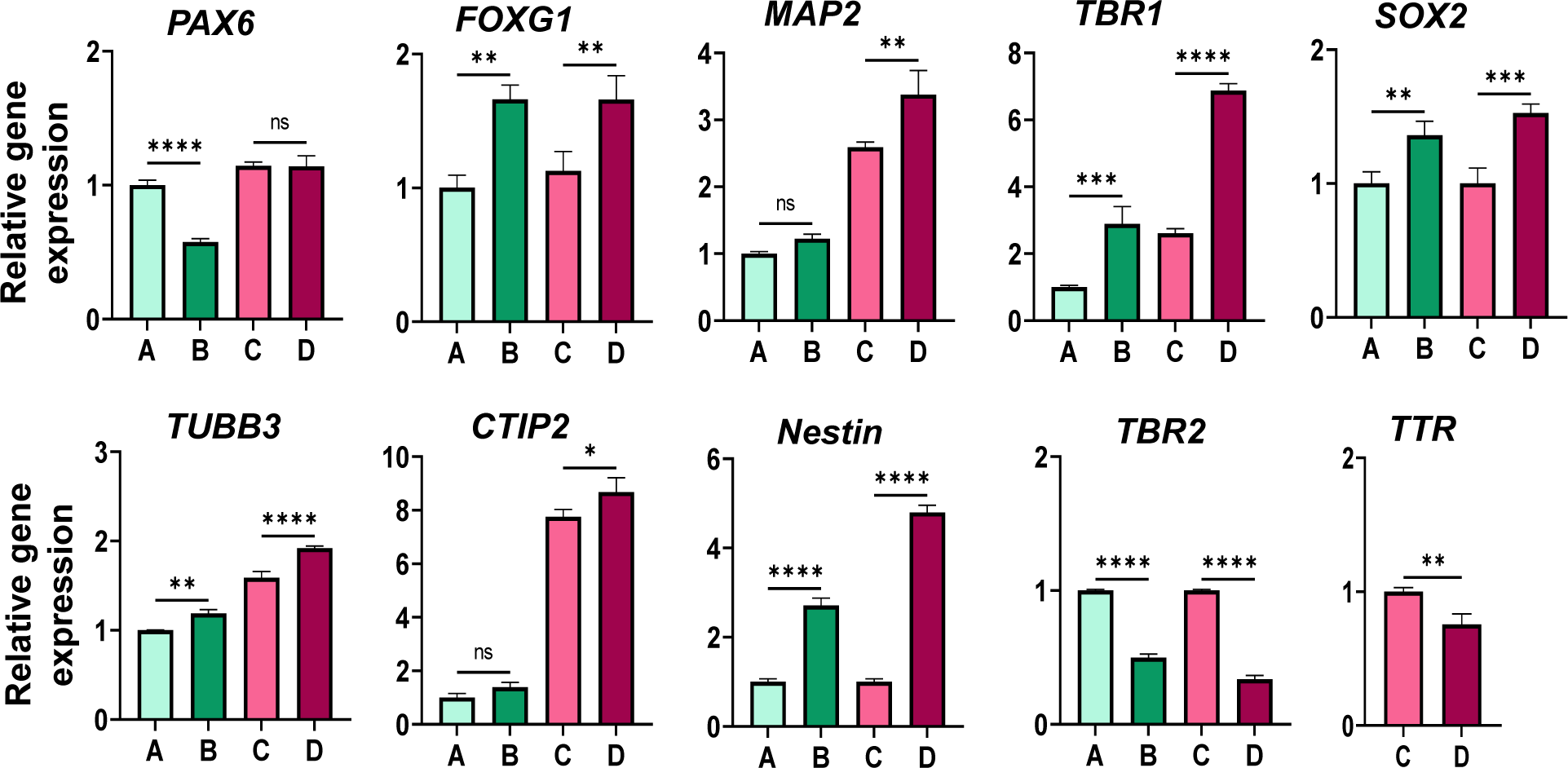
Relative neural biomarker expression in cerebral organoids: **(A)** Cultured in Matrigel domes in the 24-well plate for 16 days, **(B)** Cultured in Matrigel spots on the pillar plate for 16 days, **(C)** Cultured in Matrigel domes in the 24-well plate for 31 days, **(D)** Cultured in Matrigel spots on the pillar plate for 31 days. Neural biomarkers include *PAX6* forebrain neuroprogenitor marker, *FOXG1* forebrain marker, *MAP2* mature neuronal marker, *TBR1* preplate marker, *SOX2* proliferating neural progenitor marker, *TUBB3* neuronal cytoplasm marker, *CTIP2* deep cortical neuronal marker, *Nestin* neuronal progenitor cell marker, *TBR2* intermediate progenitor marker, and *TTR* choroid plexus marker. Statistical significance was performed by unpaired t-test. **** for p < 0.0001, *** for p < 0.001, ** for p < 0.01, * for p < 0.05, and ns = not significance (p > 0.05). n = 10 - 12 per qPCR run.

**Supplementary Figure 6.**
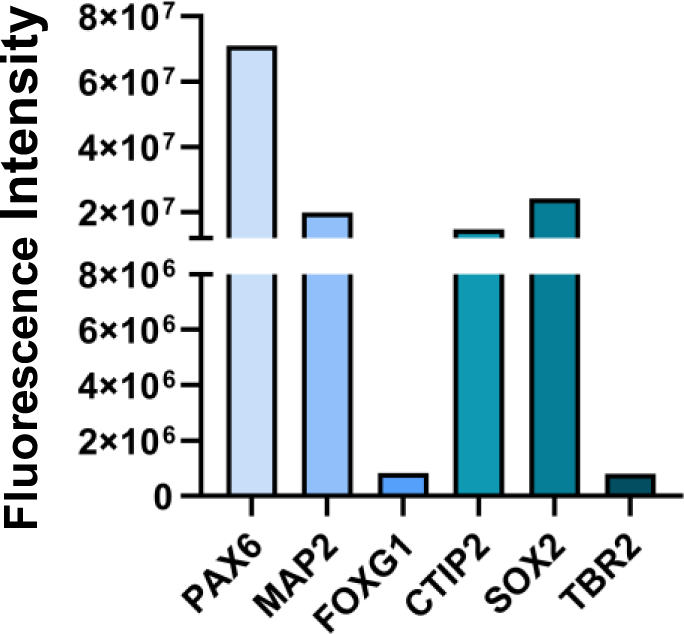
Fluorescence intensity of different biomarkers in cerebral organoids on day 31 analyzed by ImageJ software.

**Supplementary Figure 7.**
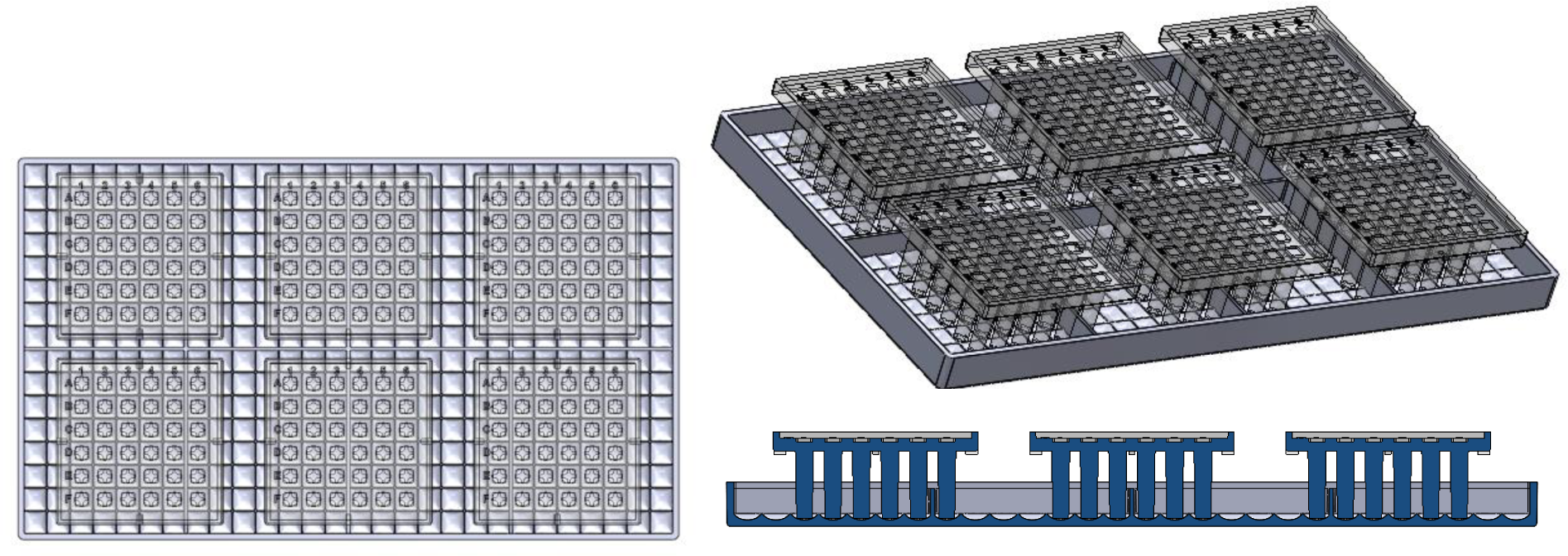
A cell loading plate for loading cells suspended in hydrogels on the pillar plate by simple one-step stamping.

